# Behavioural evidence of a humidistat: a temperature-compensating mechanism of hydroregulation

**DOI:** 10.1101/2025.01.30.635663

**Authors:** Danilo Giacometti, Glenn J. Tattersall

## Abstract

The ability to control hydration state essential for terrestrial species, especially amphibians, which are highly susceptible to dehydration. Here, we examined how temperature (17°C vs. 22°C) influenced behavioral hydroregulation in spotted salamanders (*Ambystoma maculatum*) using a laboratory humidity gradient. Salamanders defended a constant vapour pressure deficit (VPD) between temperatures by targeting higher RH at 22°C than at 17°C, possibly to compensate for increased evaporative demand at warmer temperatures. Individuals selecting higher VPDs experienced greater evaporative water loss (EWL), with larger salamanders losing more water than smaller ones after accounting for temperature. Together, these results highlight a trade-off among body size, humidity preference, and desiccation tolerance. Salamanders also rehydrated faster at 22°C than 17°C, highlighting temperature-dependent water uptake rates. Our finding that salamanders regulated a constant driving force of evaporation between temperatures suggests the ability to detect rates of EWL. Local cooling of the skin arising from evaporation is a plausible mechanism: if moist-skinned ectotherms allow for a specific, fixed decline in skin relative to core temperature, they can effectively detect the drive for evaporation. Ultimately, our study underscores the complexity of amphibian hydroregulation and emphasises the role of behaviour in maintaining hydration state.

## Introduction

The transition from aquatic to terrestrial environments was one of the most significant milestones in vertebrate evolution (Laurin, 2010). On land, water availability fluctuates both spatially and temporally, meaning that organisms face constant challenges related to the maintenance of water balance (Takei, 2015). Therefore, the ability to control hydration state is essential for terrestrial species. Effective hydroregulation may be achieved through a combination of physiological and behavioural strategies. Physiological processes include the enhancement of skin resistance to water loss (Tracy et al., 2008), the presence of waterproofing lipids in the integument (Lillywhite, 2006), or the modulation of kidney and bladder function to increase water retention (Schmidt-Nielsen and Schmidt-Nielsen, 1952; Shoemaker and Bickler, 1979). Hydroregulatory behaviours may comprise the adoption of water-conserving postures (Pough et al., 1983), aggregation (Broly et al., 2014), or burrowing (Bulova, 2002). Often, however, behavioural hydroregulation is oversimplified to the selection of humid microhabitats (but see Dezetter et al., 2023). As a result, we still have limited information about how animals perceive and process changes in habitat humidity, and how behaviour can be used to mitigate potential disruptions to the maintenance of water balance (Hillyard and Willumsen, 2011; Hillyard et al., 1998).

Among vertebrate ectotherms, amphibians in particular face elevated hydroregulatory challenges. Truly, amphibians have a relatively thin skin that is supported by blood vessels in the epidermis, which together facilitate cutaneous gas exchange (Tattersall, 2007). However, the ability to exchange gases through the skin also makes amphibians highly susceptible to evaporative water loss (EWL), especially at high temperatures (Lertzman-Lepofsky et al., 2020). The extent to which temperature impacts EWL depends on the vapour pressure deficit (VPD), which measures the drying power of air, and is determined by the difference between the saturated water vapour pressure (WVP_sat_) and the actual water vapour pressure (WVP_act_) (Anderson, 1936). Since warm temperatures typically produce high VPD, amphibians facing these conditions should suffer high evaporative demand and pay high hydroregulatory costs (Riddell et al., 2019; Riddell et al., 2024). Because of this, many have argued that thermal constraints over hydroregulation explained the recurrent pattern of low body temperatures (*T*_b_s) in amphibians, on the basis that sustaining low *T*_b_s would minimise EWL (Brattstrom, 1979). This view has been recently expanded into a framework that posits that amphibians constantly balance thermoregulation and hydroregulation to sustain performance (Greenberg and Palen, 2021; Titon and Gomes, 2017). However, while this framework has both thermosensation and hydrosensation components, our current knowledge of the former outweighs the latter (Ortega et al., 2023; Tracy et al., 2008), particularly in terms of the behaviours that contribute to the maintenance of hydration state.

One possible explanation for this knowledge gap is the difficulty in disentangling temperature and humidity effects, since most laboratory studies test amphibians in combined humidity and thermal gradients (hereafter “hydrothermal gradients”) (Bundy and Tracy, 1977; Galindo et al., 2018). While one may argue that the use of hydrothermal gradients provides a realistic scenario, any response measured in such an apparatus is by definition confounded by temperature and humidity (Mitchell and Bergmann, 2016). Because one cannot properly gauge whether the observed behaviours are innately hydroregulatory, the extensive use of hydrothermal gradients resulted in the simplistic yet prevalent view that, for amphibians, “wetter is better” (see Brattstrom, 1979). However, if one is interested in assessing humidity preference and the behaviours that underpin hydroregulation, one should test amphibians in a temperature-controlled humidity gradient (Heatwole, 1962; Shelford, 1913). For example, Spotila (Spotila, 1972) demonstrated interspecific differences in humidity preference (RH selection range: 60.4– 90% at 15°C) across 12 closely-related species of salamanders. Humidity preference was associated with microhabitat use, suggesting that salamanders were able to detect differences in air moisture content and respond in a manner that maximised hydration state (Spotila, 1972). Despite their potential, temperature-controlled humidity gradients have seldom been used in amphibian research since the 1970s (but see Chapman and Bidwell, 2023). As such, our understanding of behavioural hydroregulation remains limited.

In this study, we used a laboratory humidity gradient to assess how temperature (17°C vs. 22°C) affected behavioural hydroregulation in the spotted salamander (*Ambystoma maculatum*). We focussed on two main behavioural metrics: VPD selection and RH selection. By measuring VPD selection, one is able to obtain behavioural information based on a temperature-independent humidity gradient and gauge behavioural hydroregulation *sensu stricto* (Dezetter et al., 2023; Wolcott and Wolcott, 2001). In contrast, RH selection provides information on how changes in temperature impact the behaviours that contribute to the maintenance of hydration state (Galindo et al., 2018; Spotila, 1972). Based on this, we predicted that *A*. *maculatum* would alter RH selection to maintain a constant VPD between temperatures (Spotila, 1972). To further understand how behaviour contributes to the maintenance of hydration state, we tested for a functional trade-off among EWL, VPD selection, activity level, and temperature. We predicted that highly active salamanders that also selected high VPD would be the ones with the highest EWL; these effects should be greater at 22°C than at 17°C (Heatwole and Newby, 1972; Spight, 1968). Finally, we also tested whether rehydration rates (ReR) increased as a function of EWL and test temperature (Claussen, 1969; Spight, 1967). With this integrative approach, we aim to provide a broader understanding of the role of behaviour to the maintenance of hydration state in amphibians.

## Material and methods

### Salamander collection and husbandry

In May 2022, we collected 44 post-breeding, adult *A*. *maculatum* (22 females + 22 males) with a drift fence installed around Bat Lake, Algonquin Provincial Park, ON, Canada (45.5857°N, 78.5185°W). We recorded the sex of each salamander based on cloacal morphology (Petranka, 1998) and identified each individual based on their unique dorsal spot pattern. To transfer the salamanders to Brock University, we allocated them into plastic containers with ventilated fitted lids (34 cm x 19.6 cm x 12 cm). These containers had *Sphagnum* moss, pine needles, and water, and were placed in a transport box at ∼ 4°C to avoid overheating and dehydration during transportation.

In the lab, we housed salamanders in pairs within ventilated tanks containing coconut husk fibre, *Sphagnum* moss, PVC pipe refuges, and a water dish. These tanks were kept in a facility with controlled temperature, RH, and photoperiod. We adjusted temperature and photoperiod seasonally to simulate native habitat conditions while keeping RH constantly at 70% (Giacometti and Tattersall, 2024). We fed salamanders twice a week with mealworms, nightcrawlers, and isopods dusted in multivitamin and calcium powder. We weighed salamanders weekly to the nearest 0.01 g using an analytical scale (Mettler Toledo, model PB602-S) to monitor changes in body mass (M_b_) as a proxy of well-being; all animals maintained M_b_ throughout their period in the lab. The current study was conducted between late June and August 2024 with salamanders that had been acclimatised to summer conditions (14°C and a 14:12 light:dark cycle) for at least four weeks.

### Laboratory humidity gradient

To assess behavioural hydroregulation, we used an annular humidity gradient (total length = 120 cm, outer radius = 45 cm, inner radius = 30 cm) built by Brock University Technical Services (Figure S1). The thermally conductive floor of the gradient had copper pipes connected to a water bath (Haake™, model DC10) kept at either 17°C or 22°C. These temperatures represent, respectively, the median and maximum upper selected temperatures (*T*_sel_) by *A*. *maculatum* in the summer (Giacometti and Tattersall, 2024). The gradient chamber was divided into an inner and outer lane by a 10.5 cm tall partition attached to the gradient lid, allowing the simultaneous study of two individuals per experiment (visually isolated). To create a humidity gradient, we delivered air of different moisture content to compartments (dry, intermediate, and humid) in the gradient chamber. We created these compartments by attaching sponges (4 cm x 3cm x 9 cm) to the gradient wall every 30 cm. To ensure the sponges remained in place and allowed for salamander navigation during the experiments, we sewed the hook side of Velcro tape onto the sponges and used aquarium safe silicon sealant (Adhesive Guru, product AG310) to attach the loop side of Velcro tape to the gradient wall. The gradient lid had two venting holes every 90° relative to the horizontal axis, such that the venting holes were positioned above the centre of the dry compartment at 0°, the intermediate compartments at 90° and 270°, and the humid compartment at 180° (Figure S1).

To generate dry air at 0°, we used tubing to direct compressed air through a column of desiccant and then into the gradient at 1900 mL/min (split to direct evenly to both lanes). We generated humid air using a water vapour bubbling system connected to the venting holes at 180°. This system consisted of an 1800 mL flask filled with dechlorinated water and kept 3°C warmer than the prevailing experimental temperature. We aerated the water in the flask with an aquarium air stone (15 cm diameter) attached to tubing that received compressed air at 2100 mL/min. The pressure created inside the flask directed water vapour from the flask to the venting holes in the gradient lid (split evenly to both lanes). To generate intermediate moisture levels at 90° and 270°, we used a manifold to create a subsample line that combined air from the dry and humid lines and directed mixed air into the gradient chamber at 1000 mL/min. Since dry air is heavier than humid air (Haynes, 2016), we drew air at 250 mL/min from the dry line, and 750 mL/min from the humid line. We verified flow rates with flow meters (Aalborg Instruments & Controls Inc., model PMR1-010972) at the beginning and end of each experiment to ensure proper functioning of the humidity gradient.

With this setup, we created a stable, near-linear humidity gradient at 17°C (*R*^2^ = 0.85) and 22°C (*R*^2^ = 0.84) (Figure S2). We determined this with hygro-thermometers (Inkbird, model IBS-TH1; accuracy ± 3%) placed every 22.5° and set to record temperature and RH every 60 s for 24 h in the absence of any animals in the gradient. Since RH is a temperature-dependent measurement and therefore not readily comparable between 17°C and 22°C, we used VPD as a measure of gradient humidity between temperatures (Table S1). Following Haynes (Haynes, 2016), where *T* is temperature (°C), we calculated:

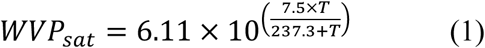

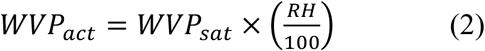

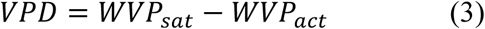

We conducted our experiments in total darkness inside a temperature-controlled room (maintained at either 17 or 22°C). We placed the humidity gradient on top of 15 cm of insulation foam board and padding so that inadvertent building vibrations would not disturb the salamanders during the experiments. We positioned an infrared illuminator (wavelength = 850 nm; TVPSii, model TP-IRBP15) and a high-resolution infrared webcam (Agama, model V-1325R) 180 cm above the centre of the gradient to record the salamanders via a time-lapse image acquisition software (HandyAVI®) set to capture an image every 30 s.

### Experimental design

We allowed the salamanders a total of 12 h inside the humidity gradient, from ∼21:00 h to ∼9:00h. Each salamander was randomly tested twice (once under each experimental temperature), with a minimum interval of 10 days between experiments. We considered the initial 3 h as the habituation period and the subsequent 9 h as the experimental period. All individuals were fasted for at least a week prior to experimentation to ensure a post-absorptive state (Secor and Boehm, 2006). We always allowed a minimum interval of 10 h between experiments after disinfecting the humidity gradient with 70% ethanol at the end of an experiment.

We always handled the salamanders using nitrile gloves. Before introducing salamanders into the gradient, we placed them individually into a container filled with 30 mL of dechlorinated water for 15 min so that water could be absorbed through the skin and animals could be introduced in a high hydration state. Then, we blotted the salamanders with paper towel, weighed them, and placed them into the gradient. We always introduced the salamanders into the gradient through the intermediate compartment, determining at random whether animals would be initially facing the dry or humid end, and whether they would be tested in the inner or outer gradient lanes. To minimise handling stress over hydroregulatory behaviours, we did not handle or manipulate the salamanders during the experiments. After finishing an experiment, we removed the salamanders from the gradient, blotted them with paper towel, and weighed them. We considered the difference in M_b_ before and after an experiment divided by the length of our experiment as a proxy for evaporative water loss (EWL; mL/h) (Ortega-Chinchilla et al., 2023). We determined ReR (mL/h) by placing the salamanders individually in a container with 30 mL of dechlorinated water for 30 min. Then, we removed the salamanders from the containers, blotted them with paper towel, and reweighed them to determine ReR based on M_b_ gained per half hour (Sherman and Stadlen, 1986). After this, we placed the salamanders back into their housing tanks.

### Data processing

We obtained a total of 1440 images for each individual over the course of each 12 h long experiment. We imported image sequences into Fiji (Schindelin et al., 2012), and for each experiment we recorded the identity (ID) and sex of the individual, as well as the test temperature (17°C or 22°C). We used the manual tracking plug-in in Fiji to track the position of the mid-body of the salamander within the gradient for each image in the sequence. As an output, we obtained a set of Cartesian (*x*, *y*) coordinates that had *x*, *y* = 0,0 as the top left of the image. We then used calculations derived from Giacometti et al. (Giacometti et al., 2021) to convert salamander body position into selected gradient humidity, which we determined by calculating median selected VPD and median selected RH.

To quantify activity in the gradient, we measured the distance moved (*d*_t_) by each individual every 30 s based on the distance defined by the arc of a circle:

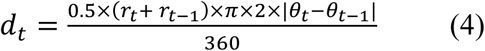

where *r* is the mean radius, and *θ* are polar coordinates between adjacent time points *t* and *t*-1 for all *n* time points. We then obtained total distance moved (*D*_t_) by taking the sum of *d*_t_:

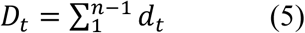

where *n* is the total number of images (*n* = 1440).

Finally, we assessed the temperature dependency of ReR, using the Van’t Hoff equation to calculate the temperature coefficient (Q_10_) for ReR between 17°C (T1) and 22°C (T2):

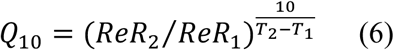

### Data analyses

We performed all analyses using R (version 4.4.1) in RStudio (version 2024-06-14) (R Core Team, 2024) considering a significance level of 0.05. We built linear mixed-effects models using the “lmer” function from the lme4 package (Bates et al., 2015), and we included ID as a random term in all models to account for repeated measures. We tested our hypotheses using data from the experimental period. We fit two individual models to test how humidity selection differed between temperatures. To this end, we considered either median VPD or RH as the response variables, experimental temperature as the predictor, and log-transformed M_b_ as a covariate. To assess how EWL differed between temperatures, we fit a model that had log-transformed EWL as the response variable, experimental temperature, median VPD, and *D*_t_ as the predictor variables, and log-transformed M_b_ as a covariate. We also evaluated how ReR differed between temperatures with a model that had log-transformed ReR as the response variable, experimental temperature and log-transformed EWL as the predictors, and log-transformed M_b_ as a covariate. For each model, we assessed residual autocorrelation with the “checkresiduals” function from the forecast package (Hyndman et al., 2020), and the “acf” and “qqnorm” functions from the stats package (R Core Team, 2024). We evaluated model fit with the “check_model” function from the performance package (Lüdecke et al., 2021), and computed marginal fixed effects through the “ggeffect” function from the ggeffects package (Lüdecke, 2018). We created figures using the ggplot2 (Wickham, 2016), Thermimage (Tattersall, 2021), and ggpubr (Kassambara and Kassambara, 2020) packages.

## Results

During the experimental period, salamanders defended a constant VPD regardless of test temperature (Table 1) (Figure 1a). This effect was mediated by M_b_, with smaller individuals selecting lower VPD than larger individuals. Importantly, salamanders avoided high VPD at both test temperatures (Figure S3). Salamanders selected different RH between temperatures (Table 1) (Figure 1b), with smaller individuals selecting higher RH than larger individuals. Regardless of temperature and activity levels, salamanders with high EWL were those that selected higher VPD and were larger (Table S2) (Figure 2). Test temperature was the only factor impacting ReR (Q_10_= 2.54 ± 2.26), with salamanders rehydrating faster at 22°C than at 17°C (Table S3) (Figure 3). Salamanders moved similar total distances within the gradient regardless of temperature (Table 2).

**Table 1.** Behavioural, body size, and physiological parameters of *Ambystoma maculatum* tested in a humidity gradient kept at either 17°C or 22°C. Values are shown as mean ± SD.

|  | 17°C<br>(N=44) | 22°C<br>(N=44) |
| --- | --- | --- |
| **Selected VPD (kPa)** |  |  |
| Mean ± SD | 0.56 ± 0.16 | 0.57 ± 0.22 |
| (min, max) | (0.27, 0.95) | (0.12, 1.06) |
| **Selected RH (%)** |  |  |
| Mean ± SD | 71.20 ± 8.31 | 78.48 ± 8.34 |
| (min, max) | (51.01, 85.88) | (60.09, 95.30) |
| **M<sub>b</sub> (g)** |  |  |
| Mean ± SD | 12.00 ± 3.91 | 12.36 ± 4.62 |
| (min, max) | (5.02, 23.70) | (4.66, 25.80) |
| **EWL (mL/h)** |  |  |
| Mean ± SD | 0.07 ± 0.05 | 0.06 ± 0.06 |
| (min, max) | (0.01, 0.19) | (0.01, 0.30) |
| **ReR (mL/h)** |  |  |
| Mean ± SD | 0.18 ± 0.13 | 0.31 ± 0.25 |
| (min, max) | (0.02, 0.56) | (0.02, 1.14) |
| **Total distance moved (m)** |  |  |
| Mean ± SD | 40.00 ± 20.80 | 40.31 ± 26.94 |
| (min, max) | (0.11, 90.62) | (0.34, 105.01) |

VPD = vapour pressure deficit, RH = relative humidity, M<sub>b</sub> = body mass, EWL = rates of evaporative water loss, ReR = rehydration rates.

**Figure 1.**
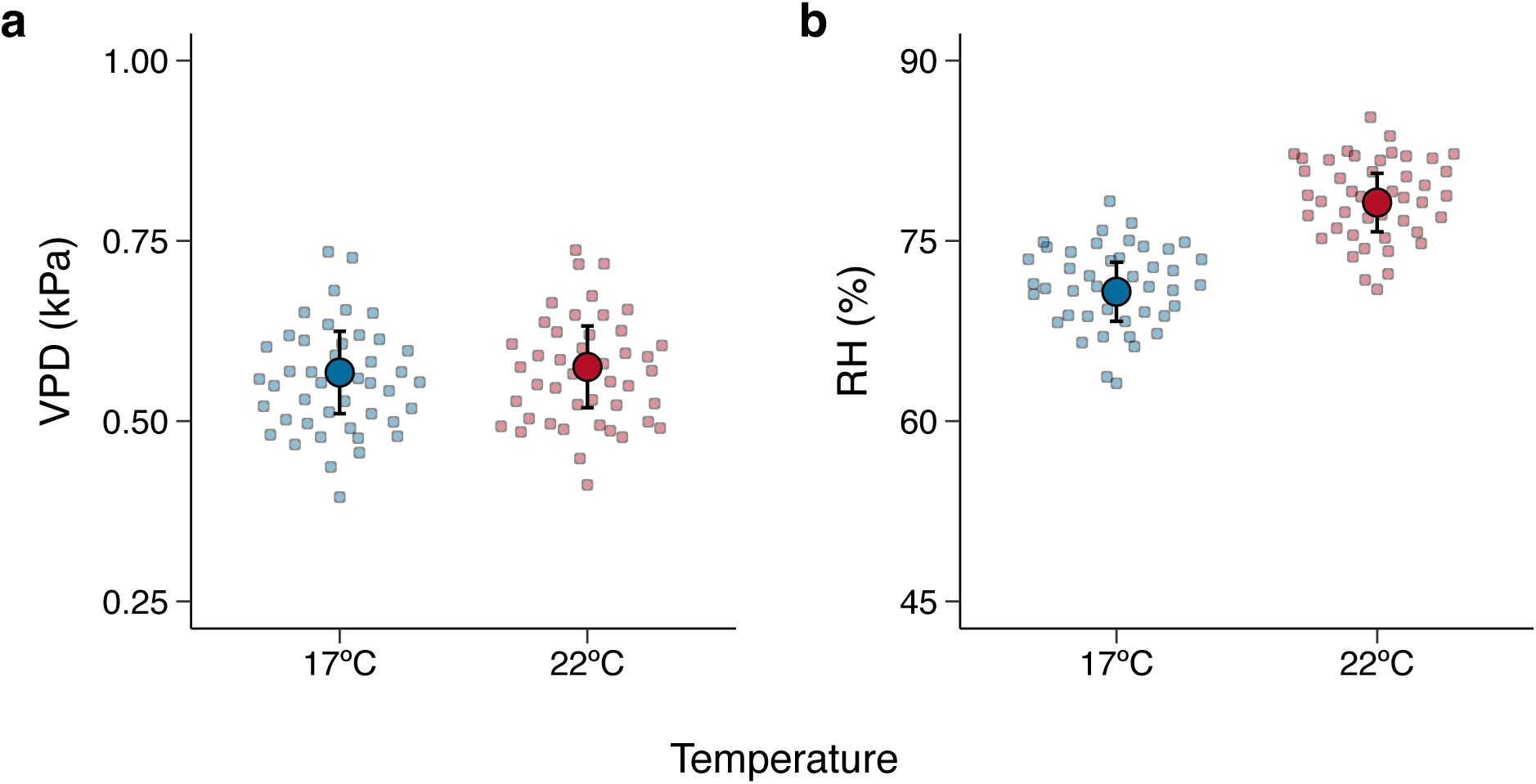
**a.** Salamanders defended a constant vapour pressure deficit (VPD) through behavioural selection of different relative humidity (RH) levels (**b**) between 17°C and 22°C. In both panels, large dots show mean marginal effects and bars indicate the corresponding 95% confidence intervals. Small dots show the mean predicted value for each individual salamander at a given temperature (17°C = blue, 22°C = red).

**Figure 2.**
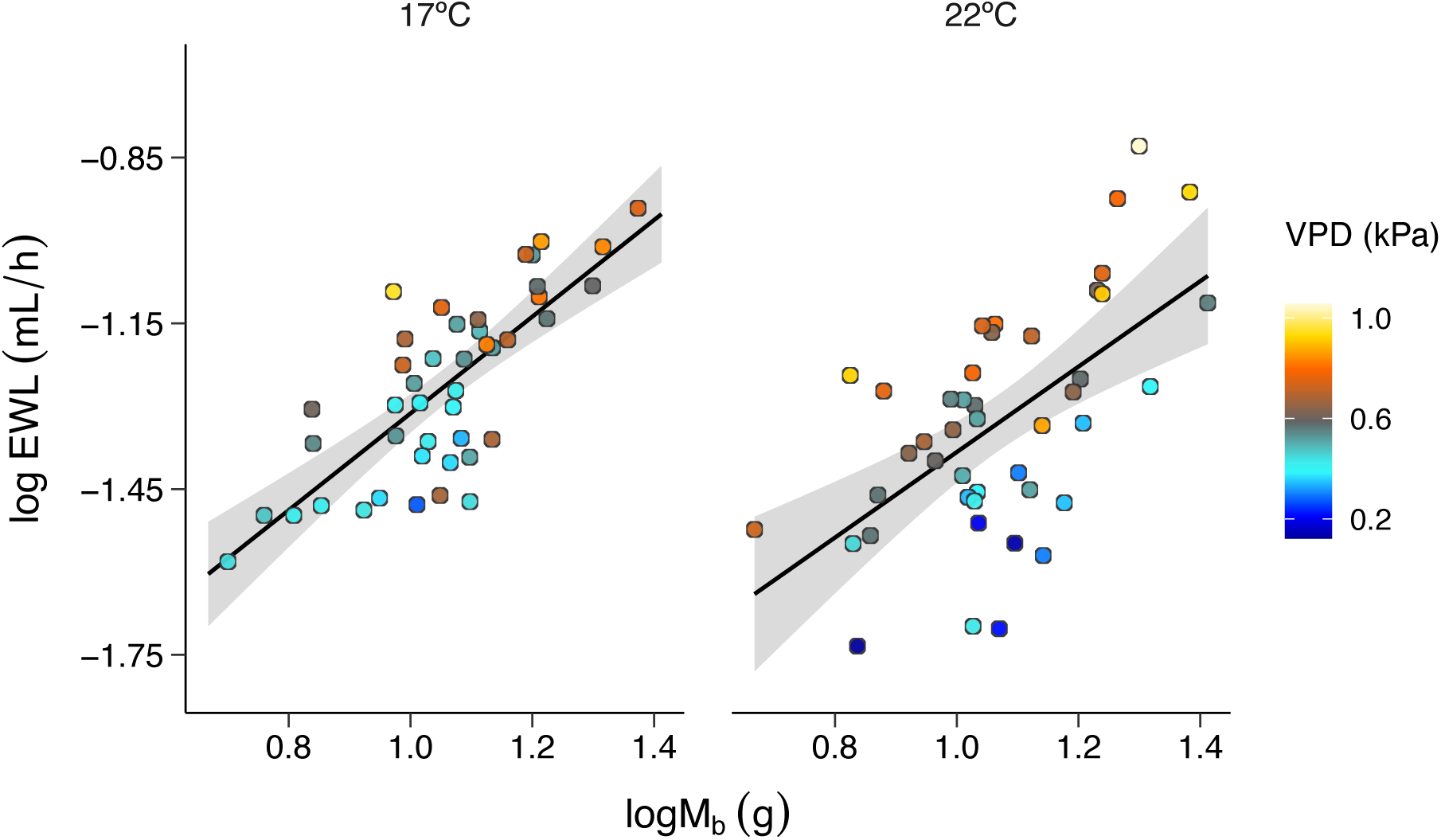
Relationship between log-transformed rates of evaporative water loss (logEWL) and log-transformed body mass (logM_b_) in *Ambystoma maculatum* tested in a humidity gradient at either 17°C or 22°C. Larger individuals had higher EWL, and individuals that selected high VPDs were also the ones with higher EWL at both temperatures. In both panels, each dot represents an individual salamander. The solid lines and shaded areas indicate the predicted relationship between logEWL and logM_b_, and the 95% confidence interval, respectively. Vapour pressure deficit (VPD) is colour-coded, with colder colours indicating wetter conditions and warmer colours indicating dryer conditions.

**Figure 3.**
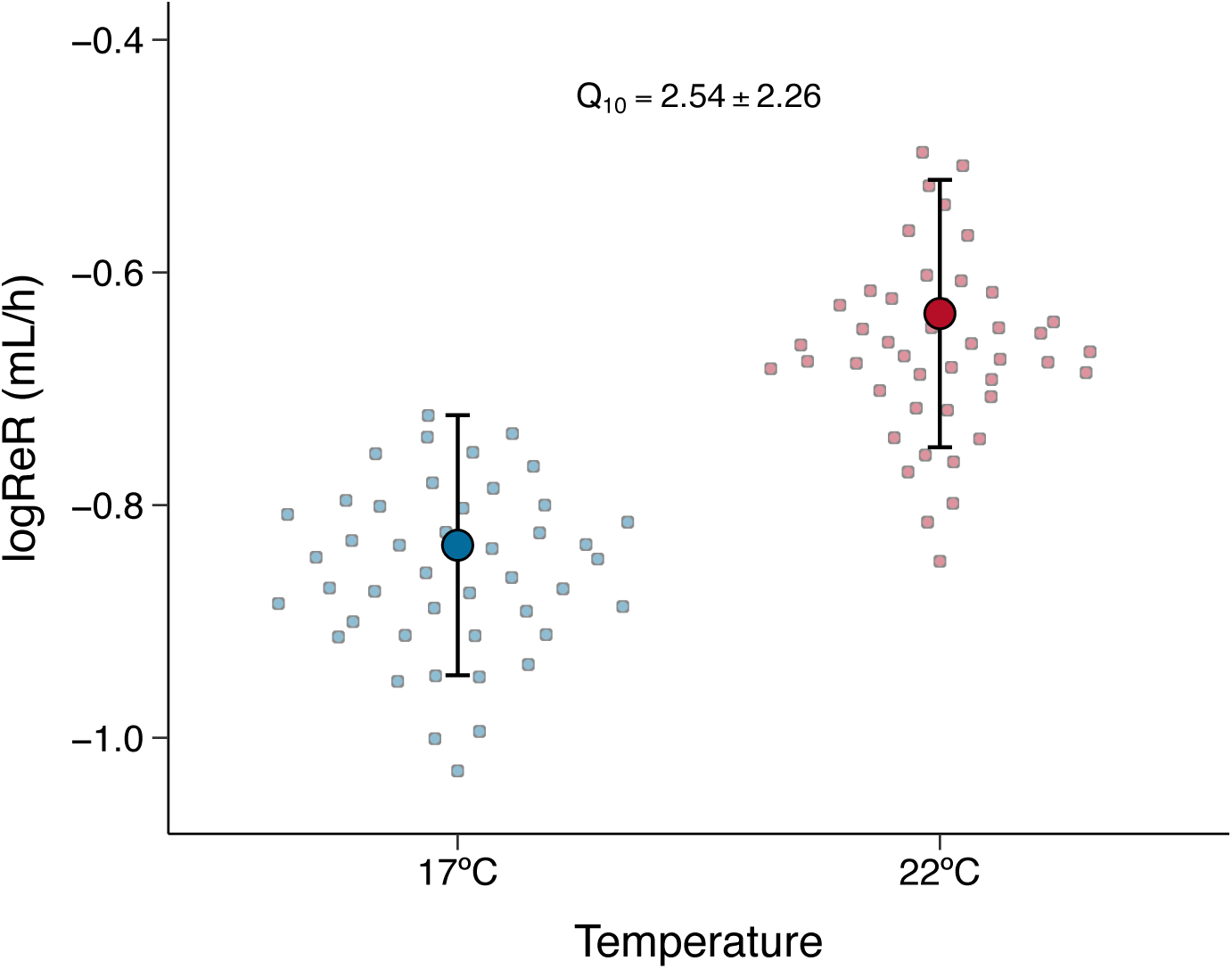
Difference in log-transformed rehydration rates (logReR) between temperatures. Salamanders rehydrated faster at 22°C than at 17°C. Large dots show mean marginal effects and bars show the corresponding 95% confidence intervals. Small dots indicate the mean predicted value for each individual salamander between temperatures (17°C in blue and 22°C in red).

**Table 2.** Parameter estimates (β), 95% confidence intervals (95% CI), and *p*-values for the models testing how humidity selection differed between temperatures in *Ambystoma maculatum* (N = 44). We considered either vapour pressure deficit (VPD) or relative humidity (RH) as response variables, temperature and log-transformed body mass (logM_b_) as predictors, and ID as a random term. Significant parameters are shown in bold.

| **VPD ~ temperature + $\log M_b$ + (1\|ID)** |  |  |  |
| --- | --- | --- | --- |
| *Predictors* | *Estimates* | *95% CI* | *p* |
| Intercept | 0.40 | 0.26 – 0.53 | **<0.001** |
| temperature (22°C) | 0.01 | -0.06 – 0.07 | 0.883 |
| $\log M_b$ | 0.01 | 0.00 – 0.02 | **0.011** |
| **Random Effects** |  |  |  |
| $\sigma^2$ | 0.03 | | |
| $\tau_{00 \text{ ID}}$ | 0.01 | | |
| ICC | 0.22 |  |  |
| Marginal $R^2$ / Conditional $R^2$ | 0.10 / 0.30 | | |
| **RH ~ temperature + $\log M_b$ + (1\|ID)** |  |  |  |
| *Predictors* | *Estimates* | *95% CI* | *p* |
| Intercept | 87.31 | 73.65 – 100.98 | **<0.001** |
| temperature (22°C) | 7.42 | 4.45 – 10.39 | **<0.001** |
| $\log M_b$ | -15.22 | -27.96 – -2.49 | **0.020** |
| **Random Effects** |  |  |  |
| $\sigma^2$ | 48.92 | | |
| $\tau_{00 \text{ ID}}$ | 16.14 | | |
| ICC | 0.25 |  |  |
| Marginal $R^2$ / Conditional $R^2$ | 0.22 / 0.42 | | |

$\sigma^2$ = residual variance; $\tau_{00 \text{ ID}}$ = individual variance; ICC = intraclass correlation coefficient.

## Discussion

We used an integrative approach to evaluate the effect of temperature on behavioural hydroregulation in *A*. *maculatum*. Our findings suggest that salamanders behaviourally defended a constant VPD by altering RH selection between temperatures. By targeting higher RH at 22°C than at 17°C, salamanders potentially compensate for increased evaporative drive at warmer temperatures. We also found evidence for a trade-off among EWL, body size, and humidity selection. Specifically, larger individuals that selected high VPDs experienced greater EWL than smaller individuals that selected low VPDs. Temperature was the only factor explaining ReR, with salamanders rehydrating faster at 22°C than at 17°C. Together, our findings underscore the crucial role of behavioural strategies in the maintenance of hydration state in amphibians.

We found that salamanders defended a constant selected VPD between temperatures. Since VPD represents the driving force for EWL (Riddell et al., 2024), our data suggest that *A*. *maculatum* can detect their rate of water loss and alter their behaviour patterns to control hydration state. To maintain homeostasis of VPD, salamanders selected higher RH at 22°C than at 17°C. The maintenance of a constant VPD across temperatures illustrates an underappreciated facet of amphibian behaviour: the ability to sense, process, and respond to moisture content in the air in a physiologically relevant manner (Shelford, 1913), and thereby maintain an effectively constant vapour pressure gradient and EWL. Under the “wetter is better” paradigm, amphibians are depicted as poor thermoregulators that sustain low *T*_b_s and always select the wettest microhabitats available to minimise dehydration (reviewed in Brattstrom, 1979). Our data challenge this view by demonstrating that *A*. *maculatum* not only showed evidence of humidity preference, but also employed behavioural adjustments to mitigate increased evaporative demand at warmer temperatures.

Our humidity preference values for *A*. *maculatum* were similar to those reported in the congener *A*. *opacum* (VPD = 0.22 kPa, or RH = 74% at 13°C) (Marangio and Anderson, 1977). Humidity preference in *Ambystoma* species, however, appears to be lower than those reported for other amphibians (Bellis, 1962; Galindo et al., 2018; Shoemaker et al., 1992). The fossorial habit of most *Ambystoma* species coupled with their relatively large body size could explain this pattern. Truly, fossorial amphibians typically bear behavioural and morphophysiological adaptations that allow for an increased independence from standing water in the environment compared to non-fossorial species (Shoemaker et al., 1992). For example, *Ceratophrys stolzmanni* uses soil humidity as a cue to adjust burrowing depth and maintain water balance (Székely et al., 2018), and *A*. *maculatum* can tolerate up to 41% of M_b_ loss through evaporation before losing function (Hall, 1922). Furthermore, our analyses showed that larger salamanders selected overall higher VPD than smaller ones at both test temperatures, suggesting that individuals with greater water content may be able to withstand greater VPDs (MacCracken and Stebbings, 2012). Small-bodied salamanders often coil to decrease the surface area available for EWL (e.g., Heatwole, 1960); however, we never observed *A*. *maculatum* assuming water-conserving postures despite the relatively wide range of body size in our study animals (M_b_ range = 4.66–25.80 g). Besides possessing a heightened desiccation tolerance (Giacometti and Tattersall, 2025) and being able to maintain hydration state behaviourally, *A*. *maculatum* may also benefit from the presence of cutaneous grooves in the venter and flanks which aid in transferring water to different parts of the body by capillarity (Lopez and Brodie, 1977; Toledo and Jared, 1993).

Our study also highlights how body size may influence behavioural hydroregulation in *A*. *maculatum*. Larger salamanders selected higher VPD and exhibited greater EWL than smaller salamanders, indicating a size-dependent trade-off between evaporation and humidity selection. Contrary to our prediction, however, the relationship between EWL and VPD was not mediated by temperature nor activity level. The allometric scaling between body size and EWL suggests that, relative to their size, smaller individuals should evaporate faster than larger individuals because small individuals have a high surface area-to-volume ratio (Jørgensen, 1997). Therefore, smaller salamanders are predicted to experience greater hydric costs than larger salamanders, and behavioural adjustments should be crucial to mitigate EWL (Riddell et al., 2024). Our results supported this notion, as evidenced by the fact that smaller *A*. *maculatum* prioritised the selection of low VPDs that culminated in overall low EWL. By contrast, larger *A*. *maculatum* selected relatively high VPDs that resulted in increased EWL. In nature, size-specific habitat use is an important behavioural strategy to minimise dehydration in aboveground-dwelling salamanders (Heatwole, 1962; Spotila, 1972). However, the extent to which size-dependent habitat selection contributes to hydroregulation in fossorial amphibians still warrants further research.

Consistent with previous work (Claussen, 1969; Spight, 1967), we found that *A*. *maculatum* rehydrated faster at 22°C than at 17°C. This thermal dependency of rehydration highlights the dual role of temperature as both a driver of EWL (Giacometti and Tattersall, 2025) and a modulator of ReR. Notably, however, we did not find a relationship between EWL and ReR in the current study. A possible reason for this lies in the fact that we allowed *A*. *maculatum* to hydroregulate at will prior to ReR assessments. Evidence from *Rhinella marina* indicates that toads may gain water at rates as fast as 30 mL/h when dehydrated to 30% of their initial M_b_ (Kosmala et al., 2020). However, water uptake is minimal when *R*. *marina* have access to water (Juarez et al., 2024). Our results back up this pattern, as rehydration in *A*. *maculatum* mostly consisted of salamanders regaining their initial M_b_. Indeed, *A*. *maculatum* lost a similar proportion of M_b_ between temperatures (6.69 ± 3.57% at 17°C and 6.01 ± 3.81% at 22°C), and thus our reported values might not represent their maximum ReRs. Despite the integumentary and vascular adaptations that favour water retention in salamanders, ReRs in this group are still lower than those of anurans (Marangio and Anderson, 1977; Spight, 1967; Toledo and Jared, 1993). Nonetheless, our calculated Q_10_ value for ReR suggests that at least some degree of temperature-dependent physiological regulation of water uptake exists in *A*. *maculatum*. The mechanisms behind such regulation are still unclear, but may involve hormonal control of skin permeability to enhance water uptake (Alvarado and Johnson, 1965), along with an increase in blood circulation at warmer temperatures (Jørgensen, 1997).

## Conclusions

The ability to maintain hydration state under varying thermal conditions is crucial to amphibians (Ortega-Chinchilla et al., 2023). By integrating behavioural and physiological perspectives, our study demonstrates the critical role of temperature-compensating behaviours to the maintenance of hydration state in *A*. *maculatum*. Our finding that salamanders maintained a constant VPD between temperatures challenges the traditional use of RH measurements to assess hydroregulation and habitat suitability in amphibians (Galindo et al., 2018). Thus, to properly understand amphibian hydroregulation, future studies should incorporate information on how behaviour is used to manage water loss, rather than rely on simple physical measurements. Although our study provides insight into the behaviours involved in hydroregulation, we still have limited information about hydrosensation compared to thermal sensation. In this context, clarifying how amphibians use water-sensing units in the skin to detect water content in the air is invaluable (Spotila, 1972). Likewise, the observation that salamanders regulate a constant driving force of evaporation under changing thermal regimes suggests a mechanism for detecting rates of EWL. Local cooling of the skin arising from evaporation (Andrade, 2016; Sinsch, 1984) is a plausible mechanism; if moist-skinned ectotherms allow for a specific, fixed decline in skin relative to core temperature, they can effectively detect the drive for evaporative water loss. Ultimately, our study underscores how behavioural adjustments are central to creating conditions conducive to the maintenance of a homeostatic hydration state in amphibians.

## Acknowledgements

We thank Ontario Parks, the Ministry of Northern Development, Mines, Natural Resources, and Forestry, the Algonquin Wildlife Research Station, Kevin Kemmish, Njal Rollinson, and Patrick Moldowan for facilitating access to Bat Lake and our study animals. We are grateful to Shawn Bukovac, Kristin Bray, Sarah Kehoe, Tom Eles, and Natasha Hearn for assistance with animal care. We thank Stephen Renda, Mitch Sillaste and Art Reimer from Brock University’s Machine Shop for building the apparatus used in our study.

## Author contributions

Danilo Giacometti and Glenn Tattersall conceived the ideas and designed methodology; Danilo Giacometti collected the data; Danilo Giacometti analysed the data with input from Glenn Tattersall; Danilo Giacometti led the writing of the manuscript. Both authors contributed critically to the drafts and gave final approval for publication.

## Data availability statement

Our data and code can be accessed from the Brock University Dataverse: https://doi.org/10.5683/SP3/IKHBTP.

