## Supplementary material for "Behavioural evidence of a humidistat: a temperature-compensating mechanism of hydroregulation": Figure S1

Supplementary tables S1, S2, and S3.

Supplementary figures S1, S2, and S3.

**Supplementary tables**

**Table S1.** Experimental conditions of the humidity gradient used in the current study (kept at either 17°C or 22°C). We measured temperature and relative humidity (RH) every 60 seconds during a 24-h period to establish baseline gradient conditions in the absence of any animals. Below, humidity is expressed both as RH and as vapour pressure deficit (VPD).

| | **17°C** | | | **22°C** | | |
|---|---|---|---|---|---|---|
| | **Dry** | **Mid** | **Wet** | **Dry** | **Mid** | **Wet** |
| **Temperature (°C)** | | | | | | |
| Mean ± SD | 17.47 ± 0.35 | 16.93 ± 0.32 | 17.33 ± 0.28 | 22.17 ± 0.34 | 21.58 ± 0.21 | 22.12 ± 0.29 |
| (min, max) | (16.42, 19.10) | (16.00, 18.47) | (16.80, 18.21) | (21.0, 22.52) | (21.10, 21.96) | (21.28, 22.39) |
| **RH (%)** | | | | | | |
| Mean ± SD | 37.56 ± 7.00 | 69.22 ± 15.09 | 93.41 ± 3.32 | 43.27 ± 5.61 | 68.50 ± 12.71 | 96.10 ± 2.04 |
| (min, max) | (26.46, 74.70) | (45.81, 96.87) | (85.73, 99.30) | (36.22, 67.27) | (50.59, 96.24) | (89.67, 99.10) |
| **VPD (kPa)** | | | | | | |
| Mean ± SD | 1.25 ± 0.14 | 0.59 ± 0.297 | 0.13 ± 0.065 | 1.51 ± 0.16 | 0.81 ± 0.33 | 0.10 ± 0.05 |
| (min, max) | (0.49, 1.50) | (0.06, 1.07) | (0.01, 0.30) | (0.81, 1.72) | (0.09, 1.30) | (0.02, 0.27) |

SD = standard deviation, min = minimum value, max = maximum value.

**Table S2.** Parameter estimates (β), 95% confidence intervals (95% CI), and *p*-values for the models testing the trade-off between evaporative water loss (EW) and vapour pressure deficit (VPD) selection between temperatures in *Ambystoma maculatum* (N = 44). We considered log-transformed EWL (logEWL) as the response variable, temperature, VPD, log-transformed body mass ($logM_b$), and total distance moved ($D_t$) as the predictors, and ID as a random term. Significant parameters are shown in bold.

| | **logEWL ~ VPD + temperature + $logM_b$ + $D_t$ + (1\|ID)** | | |
|---|---|---|---|
| *Predictors* | *Estimates* | *95% CI* | *p* |
| Intercept | -2.27 | -2.84 – -1.69 | **<0.001** |
| VPD | 0.58 | 0.15 – 1.01 | **0.009** |
| temperature (22ºC) | -0.08 | -0.21 – 0.05 | 0.208 |
| $logM_b$ | 0.64 | 0.10 – 1.18 | **0.021** |
| total distance moved | 0.00 | -0.00 – 0.00 | 0.946 |
| **Random Effects** | | | |
| $\sigma^2$ | 0.09 | | |
| $\tau_{00\ ID}$ | 0.02 | | |
| ICC | 0.20 | | |
| Marginal $R^2$ / Conditional $R^2$ | 0.21 / 0.36 | | |

$\sigma^2$ = residual variance; $\tau_{00\ ID}$ = individual variance; ICC = intraclass correlation coefficient.

**Table S3.** Parameter estimates (β), 95% confidence intervals (95% CI), and *p*-values for the models testing how rehydration rates varied as a function of evaporative water loss and temperature in *Ambystoma maculatum* (N = 44). We considered log-transformed rehydration rates (logReR) as the response variable, log-transformed rates of evaporative water loss (logEWL), temperature, and log-transformed body mass ($logM_b$) as the response variables, and ID as a random term. Significant parameters are shown in bold.

| | **logReR~ logEWL + temperature + $logM_b$ + (1\|ID)** | | |
|---|---|---|---|
| *Predictors* | *Estimates* | *95% CI* | *p* |
| Intercept | -0.83 | -1.57 – -0.09 | **0.028** |
| logEWL | 0.17 | -0.06 – 0.39 | 0.147 |
| temperature (22 ºC) | 0.20 | 0.04 – 0.36 | **0.013** |
| $logM_b$ | 0.18 | -0.38 – 0.73 | 0.526 |
| **Random Effects** | | | |
| $\sigma^2$ | 0.13 | | |
| $\tau_{00\ ID}$ | 0.00 | | |
| Observations | 88 | | |
| Marginal $R^2$ / Conditional $R^2$ | 0.10 / 0.00 | | |

$\sigma^2$ = residual variance; $\tau_{00\ ID}$ = individual variance.

### Supplementary figures

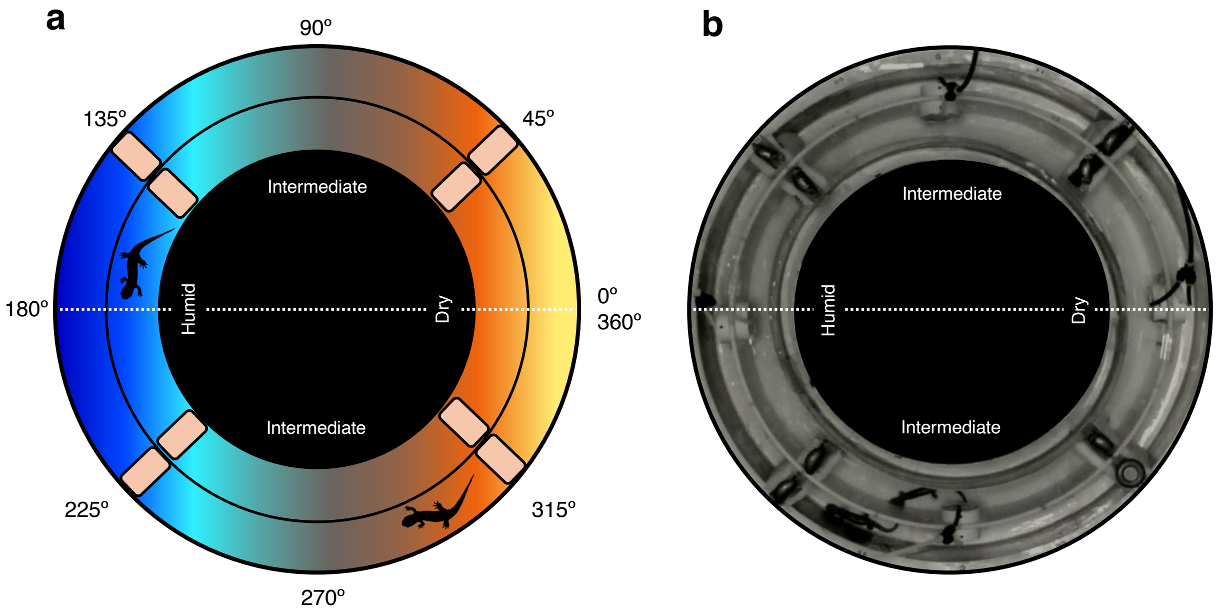


**Figure S1. a**. Schematic of the annular humidity gradient used to assess how temperature affects behavioural hydroregulation in *Ambystoma maculatum*. We affixed sponges at 45°, 135°, 225°, and 315° to maintain humidity levels within gradient compartments. Gradient humidity is colour-coded, with colder colours indicating wetter conditions and warmer colours indicating dryer conditions. **b**. Actual image of the annular humidity gradient. The hygro-thermometers at the centre of each gradient compartment recorded temperature and relative humidity every 30 s during our 12 h long experiments. Two salamanders can be seen occupying the intermediate compartment on the bottom part of the humidity gradient.

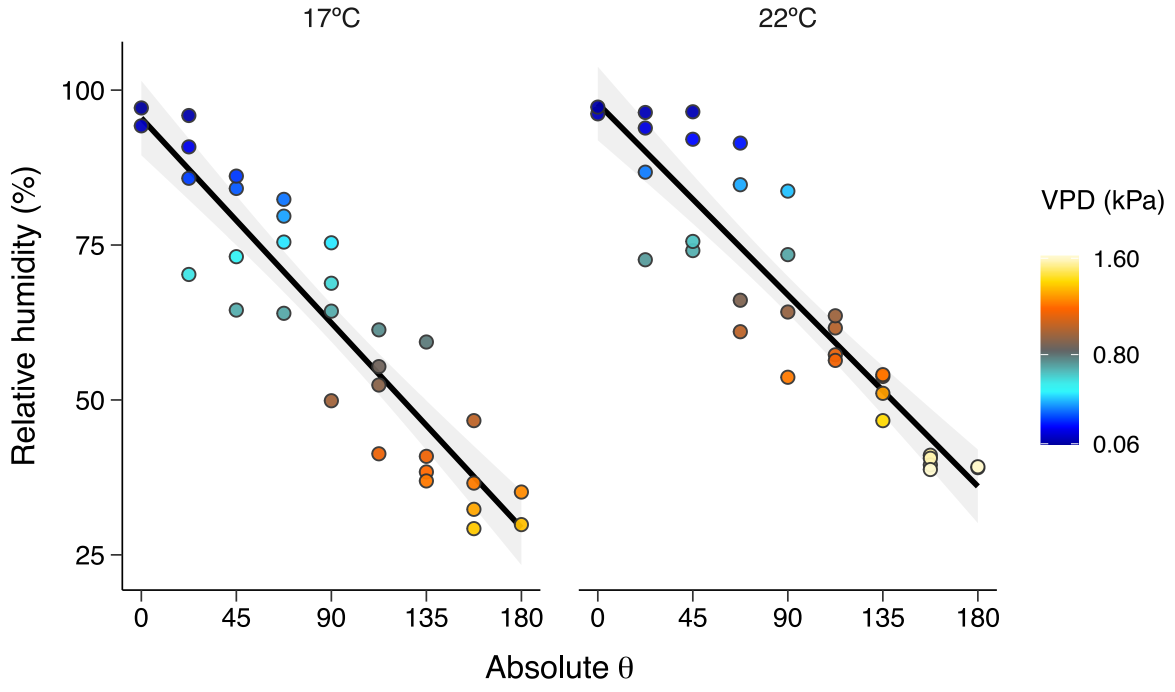


**Figure S2.** Relationship between relative humidity and polar coordinates (absolute θ) of the annular gradient used in this study (kept either at 17°C or 22°C). Vapour pressure deficit (VPD) is colour-coded, with colder colours indicating wetter conditions and warmer colours indicating dryer conditions. At 17°C, the second-order regressions describing the relationship between relative humidity and gradient position followed: $y = -1_x10^{-4}x^2 - 0.35x + 94.85$ with $R^2 = 0.85$. At 22°C, the second-order regressions describing the relationship between relative humidity and gradient position followed: $y = -3_x10^{-4}x^2 - 0.29x + 96.20$ with $R^2 = 0.84$. The solid lines and shaded areas indicate the predicted relationship between relative humidity and absolute θ, and the 95% confidence interval, respectively.

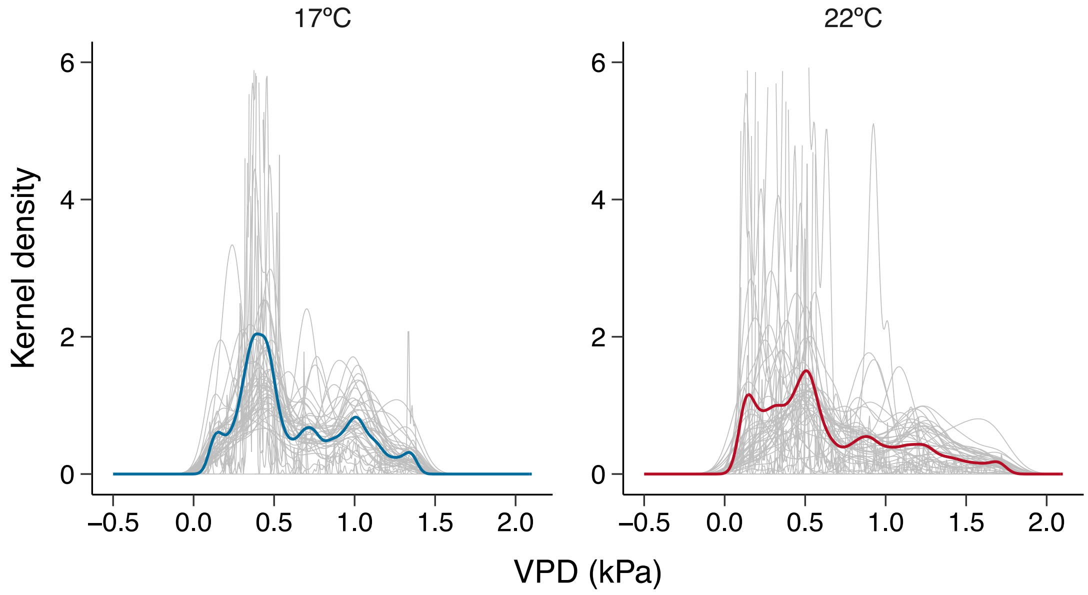


**Figure S3.** Kernel density estimates of median selected VPD in *Ambystoma maculatum*. In both panels, grey lines show the individual distribution of VPD selection, and the colour-coded lines show the median VPD distribution considering all individuals tested at a given temperature. Values above 6 on the y-axis depict salamanders that selected a specific VPD for most of the experiment. The x-axis was expanded to negative values to include the tail of the kernel density estimates.
